# SaLSa: a combinatory approach of semi-automatic labeling and long short-term memory to classify behavioral syllables

**DOI:** 10.1101/2023.04.05.535796

**Authors:** Shuzo Sakata

**Author notes:** Correspondence should be address to Shuzo Sakata. Author Contributions: SS designed research, performed research and wrote the paper. Funding Sources: MRC (MR/V033964/1) and Horizon2020-ICT (DEEPER, 101016787) to SS.

## Abstract

Accurately and quantitatively describing mouse behavior is an important area. Although advances in machine learning have made it possible to track their behaviors accurately, reliable classification of behavioral sequences or syllables remains a challenge. In this study, we present a novel machine learning approach, called SaLSa (a combination of semi-automatic labeling and long short-term memory-based classification), to classify behavioral syllables of mice exploring an open field. This approach consists of two major steps: first, after tracking multiple body parts, spatial and temporal features of their egocentric coordinates are extracted. A fully automated unsupervised process identifies candidates for behavioral syllables, followed by manual labeling of behavioral syllables using a graphical user interface. Second, a long short-term memory (LSTM) classifier is trained with the labeled data. We found that the classification performance was marked over 97%. It provides a performance equivalent to a state-of-the-art model while classifying some of the syllables. We applied this approach to examine how hyperactivity in a mouse model of Alzheimer’s disease (AD) develops with age. When the proportion of each behavioral syllable was compared between genotypes and sexes, we found that the characteristic hyper-locomotion of female AD mice emerges between 4 and 8 months. In contrast, age-related reduction in rearing is common regardless of genotype and sex. Overall, SaLSa enables detailed characterization of mouse behavior.

**Significance Statement:** Describing complex animal behavior is a challenge. Here, we developed an open-source, combinatory approach to behavioral syllable classification, called SaLSa (a combination of **s**emi- **a**utomatic labeling and **l**ong **s**hort-term memory-based cl**a**ssification). In order to classify behavioral syllables, this approach combines multiple machine learning methods to label video frames semi-automatically and train a deep learning model. To demonstrate SaLSa’s versatility, we monitored the exploratory behavior of an Alzheimer’s disease mouse model and delineated their complex behaviors. We found that female Alzheimer’s mice become hyperactive in the sense that their locomotion behavior, but not other active behaviors, appear more frequently than controls and even male Alzheimer’s mice as they age. SaLSa offers a toolkit to analyze complex behaviors.

## Introduction

In modern neuroscience, a goal is to understand the relationship between behavior and neural ensembles. However, accurately and quantitatively describing complex behavior remains challenging. In the past, mouse behaviors have often been assessed using simple, subjective criteria in a series of behavioral tests. However, advances in machine learning have changed the field (Mathis et al., 2020; Pereira et al., 2020; Luxem et al., 2023).

To describe mouse behaviors, several steps are required. First, behaviors are video-monitored. The state-of-the-art is to utilize a depth camera (Wiltschko et al., 2020) or multiple cameras (Dunn et al., 2021; Schneider et al., 2022). While some experiments require video recording from the bottom to monitor limb movement (Pereira et al., 2019; Bohnslav et al., 2021; Luxem et al., 2022), most experiments still utilize videos taken from the top. Second, the movement of the mouse’s body parts is tracked frame-by-frame to estimate animal posture. Deep learning-based algorithms have been widely adopted for this purpose (Mathis et al., 2018; Pereira et al., 2019). The final step is to identify and classify distinct behavioral sequences or syllables. While a variety of approaches have been developed over the past decade (Kabra et al., 2013; Hsu and Yttri, 2021; Luxem et al., 2023), this final step is still challenging.

There are two broad categories of methods used to classify behavioral syllables. The first category is a top-down approach, which involves pre-defining a set of rules to identify behavioral syllables or applying supervised machine learning to classify them (Kabra et al., 2013; Segalin et al., 2021; Harris et al., 2023). The second category is a bottom-up approach, which involves analyzing data patterns using unsupervised classification algorithms (Dunn et al., 2021; Hsu and Yttri, 2021; Gabriel et al., 2022; Jia et al., 2022; Luxem et al., 2022; Weinreb et al., 2023). Either approach has its own advantages and disadvantages. For example, although the top-down approach provides interpretable outcomes by definition, it can be laborious to prepare labeled datasets for model training. Conversely, while the bottom-up approach is less unbiased, setting optimal parameters and providing each syllable to an interpretable behavioral label can be a non-trivial task.

Here we hypothesize that the long short-term memory (LSTM) model (Hochreiter and Schmidhuber, 1997) can be adopted for this purpose due to the following reasons: first, behavioral sequences may contain contextual, long-term dependencies (Musall et al., 2019; Issa et al., 2020). Second, the LSTM model is designed to handle time series data with long-term dependencies and has shown promising results in other fields (Vinyals et al., 2019; Van Houdt et al., 2020). While this model has not been adopted to classify behavioral syllables, it may be a suitable method for addressing the challenges associated with these sequences. Here we introduce SaLSa (a combination of **s**emi- **a**utomatic labeling and **LS**TM-based cl**a**ssification) (**Figure 1**). This is a combinatorial approach to creating labeled data semi-automatically and training an LSTM classifier. To demonstrate the capability of this approach, we examine how behavioral abnormalities in an Alzheimer’s mouse model emerge during aging.

**Figure 1.**
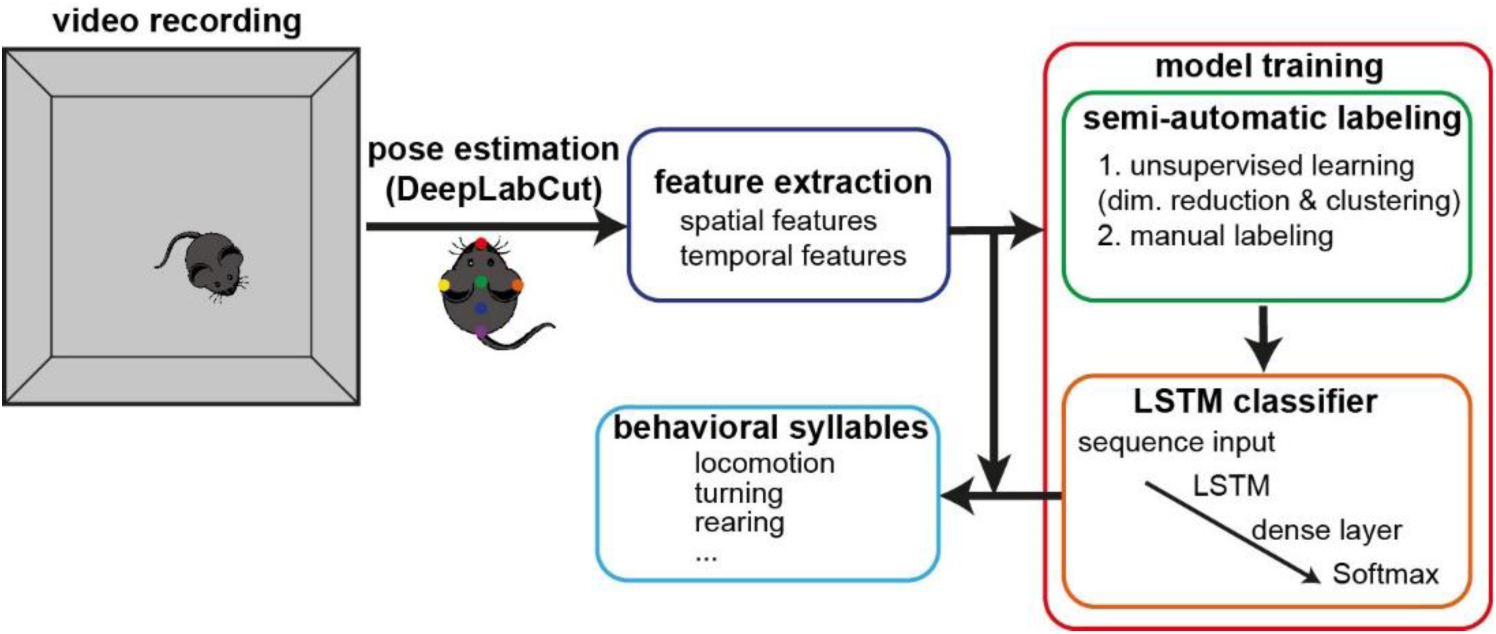
SaLSa. After recording videos and processing them with DeepLabCut (“pose estimation”), spatial and temporal features are extracted from the egocentric coordinates of tracked body parts (feature extraction). Based on a set of videos, an LSTM classifier is trained (model training). This component consists of two parts: first, through fully automated unsupervised processes including dimensionality reduction and clustering, behavioral syllable candidates are identified. Using a graphical user interface, the identified candidates are manually labeled (semi-automatic labeling). For the training and evaluation of an LSTM classifier, labeled data will be used. Once the classifier is trained, the entire sequence of extracted features is processed to classify behavioral syllables.

## Materials and Methods

### Animals

All animal experiments were performed in accordance with the United Kingdom Animals (Scientific Procedures) Act of 1986 Home Office regulations and approved by the University of Strathclyde Animal Welfare and Ethical Review Body and the Home Office (PPL0688994). 5xFAD mice (JAX006554) (Oakley et al., 2006) were bred with wild-type (WT) mice on the C57BL/6J background (>F10). All genotyping was performed by Transnetyx using real-time PCR. 43 mice (13 5xFAD male; 11 5xFAD female; 9 WT male; 10 WT female; age range: 1.3 - 9.4 months old) were used. WT mice were littermates. They had ad libitum access to food and water. The animals were housed with their littermates on a 12-h light/dark cycle. All behavioral experiments were performed during the first quarter of the light period (Zeitgeber time 2 - 3).

### Video monitoring of exploratory behaviors

The behavioral arena was an acrylic transparent box (40 cm x 40 cm x 40 cm, Displaypro). Paper sheets covered the outside of all side walls and landmark images (e.g., large stripes and crosses) were placed on two walls. Four boxes were placed closely on white tables (Lack Side Table, IKEA). A webcam (C900, NULAXY) was set over the boxes. The pixel resolution was 0.74 pixels/mm at the bottom of the arena. The video was captured at 25 fps by a custom-written program (LabVIEW, National Instruments). For each behavioral session, an animal was placed in the center of the arena and allowed to explore it for 20 min. After the test, the arena was cleaned with 70% ethanol. Only one session was held per day.

### Pose estimation

Every video file was cropped into four videos, each containing one box. This was done with custom-written MATLAB code. A DeepLabCut model (Mathis et al., 2018; Lauer et al., 2022) was trained: four videos were chosen from one behavioral session. 50 frames from each video were manually labeled. The labeled body parts consisted of (1) nose, (2) head, (3) left ear, (4) right ear, (5) neck, (6) anterior back (back 1), (7) posterior back (back 2), (8) tail base, (9) mid-tail, and (10) tail tip (**Figure 2A**). The number of training iterations was 410000. All videos were processed with this trained model. Filtered data was used. Because the tail shape did not reflect behavioral syllables, the mid-tail and tail tip were excluded from further analysis in this study. Thus, 8 body parts were analyzed.

**Figure 2.**
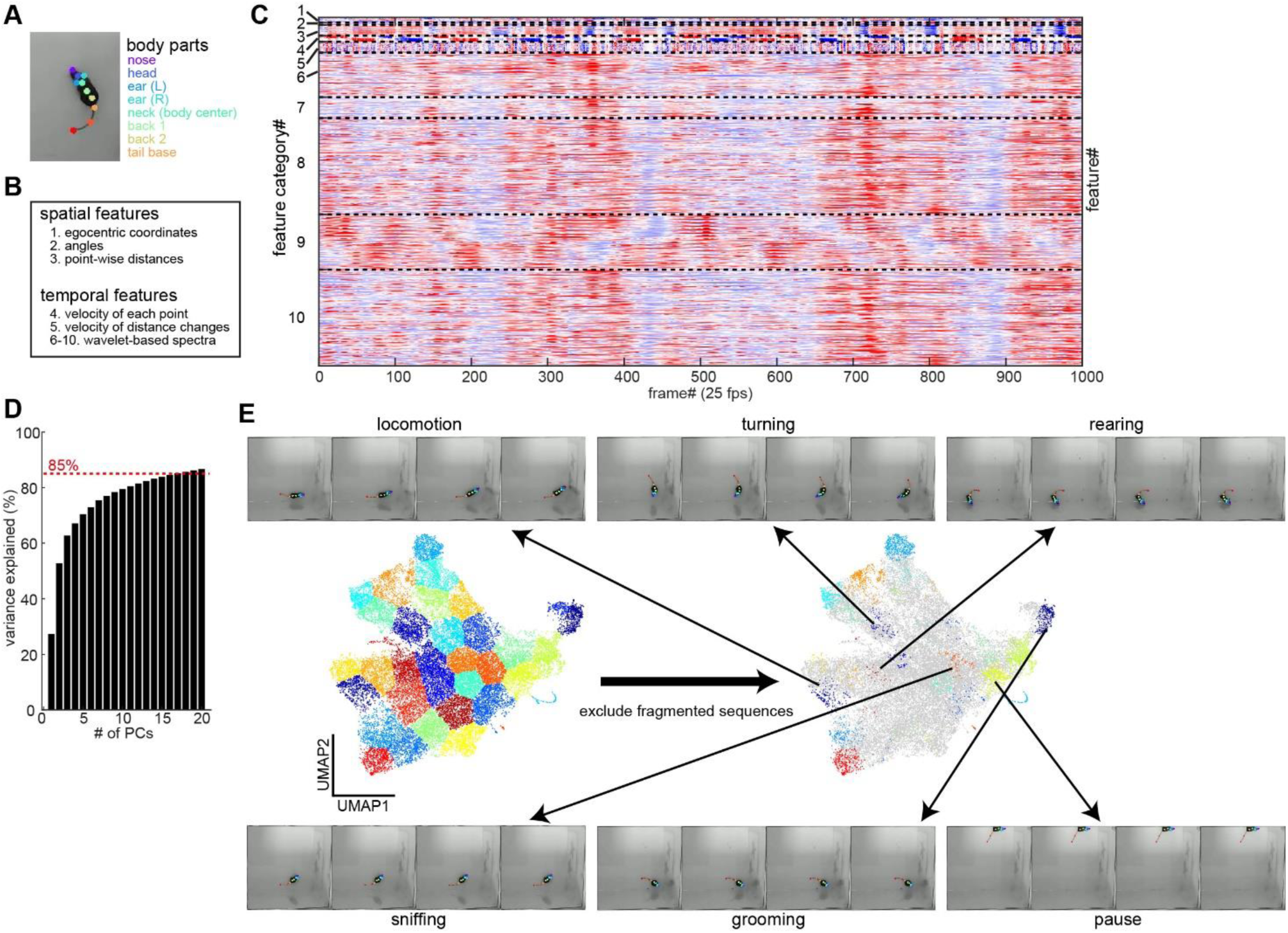
Semi-automatic labeling. (**A**) A frame with tracked body parts. Although 10 body parts were tracked, the mid-tail and tail tip were excluded for quantitative analyses. (**B**) The categories of extracted features. (**C**) A chunk of normalized feature values. Each feature was Z-scored for a visualization purpose. Dotted lines separate different feature categories. The feature category number corresponds to the number indicated in (**B**). (**D**) Cumulative distribution of explained variance across principal components (PCs). The threshold for including PCs was set at 85% variance explained. (**E**) UMAP representation of the entire video sequences and example frame sequences of labeled behavioral syllables. Data reduced by principal component analysis (PCA) was further reduced to 2 dimensions by UMAP. A spectral clustering algorithm was applied. By removing fragmented (<0.25 s) sequences, each cluster was manually annotated. The example sequences indicate a down-sampled sequence of each labeled behavioral syllable. Please note that although the mid-tail and tail tip were shown in sample frames, because the tail shape did not reflect behavioral syllables, the mid-tail and tail tip were excluded from all quantitative analyses including the UMAP analysis.

### Feature extraction

All analyses were implemented in MATLAB (https://github.com/Sakata-Lab/SaLSa). All coordinates were converted into egocentric coordinates as the neck was set as the body center and the nose-neck axis was set as the body-center axis. After conversion, comprehensive features were derived. Features can be categorized into spatial and temporal features.

The spatial features were as follows: (1) the relative coordinates across body parts, (2) the relative angle of each body part relative to the body-center axis, and (3) the distance between body parts. The temporal features were as follows: (1) the frame-by-frame velocity of each body part, (2) the frame-by-frame velocity of the distance between body parts, (3) the spectrotemporal characteristics of the relative coordinates across body parts, (4) the spectrotemporal characteristics of the relative angle of each body part, (5) the spectrotemporal characteristics of the frame-by-frame velocity of each body part, (6) the spectrotemporal characteristics of the distance between body parts, and (7) the spectrotemporal characteristics of the frame-by-frame velocity of the distance between body parts. To compute spectrotemporal characteristics, we applied a wavelet transformation (*cwt* function in MATLAB). The extracted frequency components were evenly spaced in the wavelet frequency domain: 0.62, 0.88, 1.25, 1.77, 2.50, 3.54, 5.01, 7.09, and 10.0 Hz. Overall, 910 features were extracted.

### Unsupervised processes

The following steps automatically identified candidates for behavioral syllables based on the extracted features. This step consisted of the following sub-steps: the first step was dimensionality reduction. Principal component analysis (PCA) was performed to reduce the dimension. PCs that explained >85 % variance were used for uniform manifold approximation and projection (UMAP) (https://www.mathworks.com/matlabcentral/fileexchange/71902). The parameters, *min_dist*, and *n_neighbors*, were set to 0.3 and 5, respectively. The second step was clustering. In the 2D UMAP space, spectral clustering was performed. The number of clusters was optimized to between 35 and 50 clusters using Calinski-Harabasz Criterion (Calinski and Harabasz, 1974). Because the main aim was to extract clusters of certain behavioral syllables with less contamination, we intentionally over-clustered the data. After clustering, each cluster contains a number of sequences of candidate behavioral syllables. The final step is a post-processing. Because short sequences were hard to label manually, fragmented (<0.25 s) sequences were excluded from labeling. This step drastically reduced sequences and clusters. After this processing, a snippet for each cluster was created for manual labeling.

### Manual labeling

A custom-written graphical user interface (GUI) was used to label each snippet. According to the initial evaluation, the following six behavioral syllables were often observed: (1) locomotion: walking/running behavior toward one direction, (2) turning: turning behavior at the same position, (3) rearing: rearing behavior toward a wall or at the center of the arena, (4) sniffing: rhythmic sniffing behavior, (5) grooming: grooming behavior, (6) pause: behavior standing still at the same position. Each snippet was labeled as one of these six syllables or miscellaneous where more than two behavioral syllables were contaminated, or initially unrecognized behaviors (such as jumping) were observed. Because this labeling procedure was for the training of an LSTM classifier, we took a conservative approach so that snippets labeled as one of 6 syllables can be less contaminated by other syllables. Out of 43 videos, 29 were manually labeled. 19 videos were used for LSTM model training. 10 additional videos were used for the model performance assessment.

### LSTM training and classification

The LSTM classifier was comprised of (1) an input layer, (2) an LSTM layer, (3) a fully connected layer, (4) a Softmax layer, and (5) a classification layer. The input layer consisted of 910 units as 910 features were extracted from each video (see above). The LSTM layer contained 200 hidden units. The classifier aimed to classify 6 behavioral syllables. For training, the ADAM optimizer was used with the L_2_ norm regularization method. The relationship between three parameters (the gradient threshold, the maximum number of epochs, and the number of hidden units) and evaluation accuracy was systematically assessed as part of optimization. In this study, the gradient threshold was set to 1 and the maximum number of epochs was set to 60. For training, 80% of labeled data was used whereas the remaining labeled data was used for evaluation. The model with the best validation performance was used for further analysis. Once the classifier was trained, all videos were processed to determine behavioral syllables frame-by-frame.

### Classification performance assessment

To evaluate the LSTM classifier’s performance, additional 10 videos were used. For each behavioral syllable, a receiver operating characteristic (ROC) curve and the area under the curve (AUC) were computed.

### Benchmark testing

To compare SaLSa’s performance with an existing model, keypoint-MoSeq (Weinreb et al., 2023) was chosen for the following reasons: first, it overperformed other major models, such as B-SOiD (Hsu and Yttri, 2021) and VAME (Luxem et al., 2022). Second, the implementation is straightforward with a limited set of parameters to explore. For training of the keypoint-MoSeq (kpms) model, we took 29 videos used for training and performance assessment for SaLSa since manually labeled data was available. The hyperparameter, *kappa*, was set to 9e4, and training iterations were set to 250 times. The frequency cutoff for behavioral syllables was set to 0.5%. These sub-threshold syllables were merged as a single syllable for post-hoc analysis. The trained model was used to analyze all these 29 videos to compare the performance of both SaLSa and kpms models with each other. For more direct comparisons between the two models, kpms’ syllables were re-assigned based on the comparison data with the manually labeled data: each original syllable was re-assigned as the most presented syllable of six pre-defined syllables (i.e., locomotion, turning, rearing, sniffing, grooming, and pause).

As an additional comparison, a multiclass support vector machine was trained to classify 6 syllables with MATLAB’s *fitcecoc* function, and performance was assessed. Similar to the LSTM model, 19 labeled videos were used for training whereas the remaining 10 videos were used for performance assessment.

### Quantification of behavioral syllables

To quantify the features of each behavioral syllable, three metrics were computed. First, speed (cm/s) was calculated in each episode. This was done by measuring the nose travel distance across frames. Second, the turning angle (°/s) was computed for each episode. This was done by calculating the cumulative angle change between the nose and tail base across frames. Finally, “compactness” was defined as the mean distance between two body parts. Smaller compactness results from the squeezed body pose.

### Statistical analysis

All statistical analyses were performed in MATLAB (version 2022a). In **Figure 5**, the Shapiro-Wilk test was performed with a Bonferroni correction to check normality. Then, a one-way analysis of variance (ANOVA) with a post-hoc Tukey HSD test was carried out. In **Figure 6**, after performing the Shapiro-Wilk test, an analysis of covariance (ANCOVA) with a post-hoc Tukey HSD test was carried out. A *p*-value less than 0.05 was considered significant. Otherwise stated, the error bars represent SEM.

**Figure 5.**
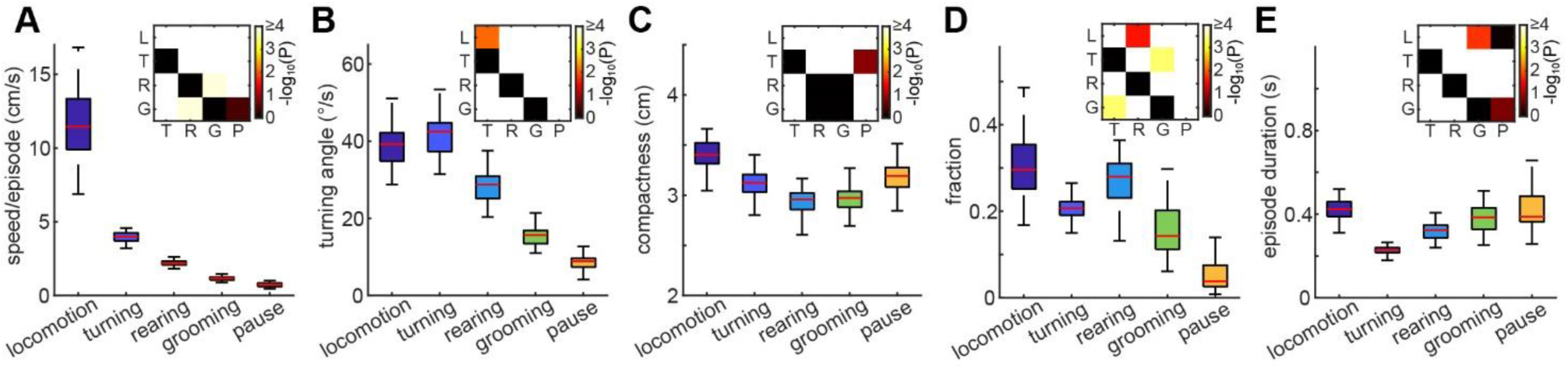
Quantification of behavioral syllables. (**A**) The median speed per episode of each video across behavioral syllables. (**B**) The median turning angle per episode of each video across behavioral syllables. (**C**) The median compactness per episode of each video across behavioral syllables. (**D**) The fraction of each behavioral syllable. (**E**) The average episode duration of each video across behavioral syllables. *inset*, *p*-values of post-hoc pair-wise comparisons (n = 43, two- way ANOVA with post-hoc Tukey HSD test). L, locomotion; T, turning; R, rearing; G, grooming; P, pause.

**Figure 6.**
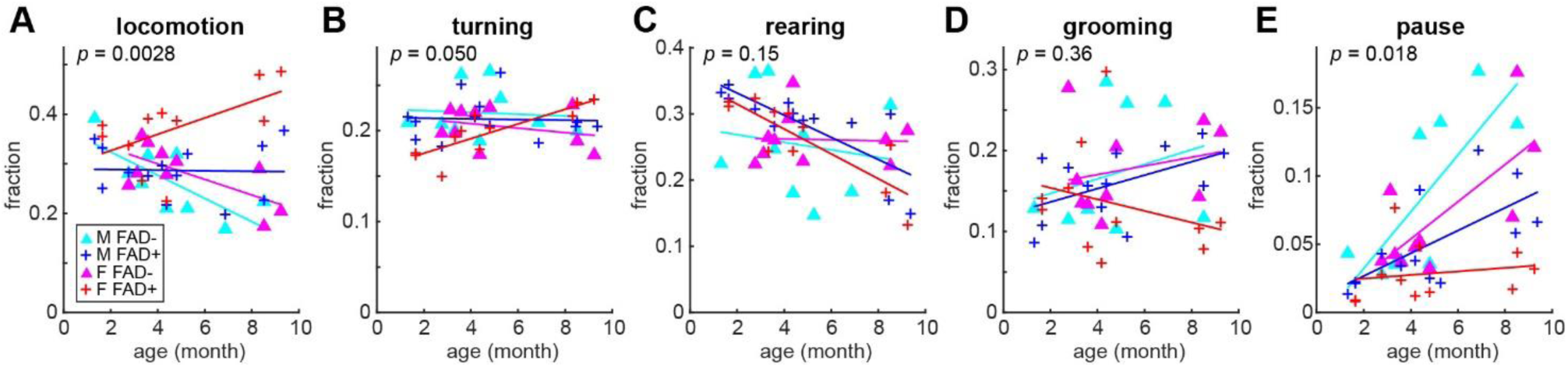
Age-dependent effects of genotype and sex on behavioral contents in 5xFAD mice. The fraction of each behavioral syllable as a function of age. Behavioral syllables are (**A**) locomotion, (**B**) turning, (**C**) rearing, (**D**) grooming, and (**E**) pause. *P*-values of ANCOVA are shown.

### Code availability

MATLAB implementations of SaLSa are publicly available (https://github.com/Sakata-Lab/SaLSa).

## Results

### SaLSa (a combination of semi-automatic labeling and LSTM-based classification)

The general workflow was as follows (**Figure 1**) (see also Materials and Methods): (1) video recording, (2) body part tracking, (3) feature extraction, (4) classifier training, and (5) classification. The step of classifier training consists of two major components (**Figure 1**): first, labeled data is prepared semi-automatically. This component starts with an unsupervised approach to automatically identify behavioral syllable candidates. This facilitates the subsequent manual labeling step. The second component is the training of an LSTM-based classifier. Using the labeled data, an LSTM classifier is trained to classify sequential data. We call this integrative approach “SaLSa” (a combination of **s**emi-**a**utomatic labeling and **LS**TM-based cl**a**ssification).

To deploy SaLSa, we began by collecting 43 videos where mice explored an open arena for 20 minutes. As described below, 5xFAD mice and their littermates of both sexes (1.3 – 9.4 months old) were used. To track multiple body parts (**Figure 2A**), a DeepLabCut (DLC) model was trained and all videos were analyzed. The DLC model test error was 1.84 pixels. Because the tail shape did not reflect behavioral syllables, the mid-tail and tail tip were excluded from further analysis in this study. Thus, 8 body parts were analyzed.

After converting all body part coordinates into egocentric coordinates as the neck was defined as the body center, we extracted 910 features with respect to spatial and temporal features (see Materials and Methods) (**Figure 2B**). **Figure 2C** shows an example sequence of all features. There were notifiable patterns across frames, implying distinct behavioral syllables.

To identify syllable candidates, we adopted unsupervised methods, including principal component analysis (PCA), uniform manifold approximation and projection (UMAP), and spectral clustering (**Figures 2D and 2E**). First, 910-dimensional data was reduced into ∼15 dimensions by using PCA so that >85% variance could be explained (**Figure 2D**). UMAP was adopted for further reduction to 2 dimensions. In this 2-dimensional UMAP space, around 40 clusters were separated using the spectral clustering algorithm. Because we aimed to identify clusters, each containing frames related to a certain behavioral syllable with minimum contamination of other syllables, we intentionally set the number of clusters high (between 35 and 50) (**Figure 2E**).

After removing fragmented, short (<0.25 s) sequences, each cluster was manually labeled as one of the following categories: (1) locomotion, (2) turning, (3) rearing, (4) sniffing, (5) grooming, (6) pause, and (7) miscellaneous (**Figure 2E**). Out of 43 videos, 29 videos were manually labeled: 19 were used for classifier training whereas 10 were used for classifier evaluation in this study.

### Classification Performance

From 19 videos, 38202 frames were labeled as one of six behavioral syllables (**Figure 3A**). While most of the frames were labeled as locomotion, less than 1% of the frames were labeled as sniffing. Despite this uneven distribution of labeled frames, the evaluation performance of the trained LSTM classifier was 97.9%. A range of parameters (i.e., the gradient threshold, the maximum number of epochs, and the number of hidden units) were explored to confirm similar evaluation performance (**Extended Data Figure 3-1**).

**Figure 3.**
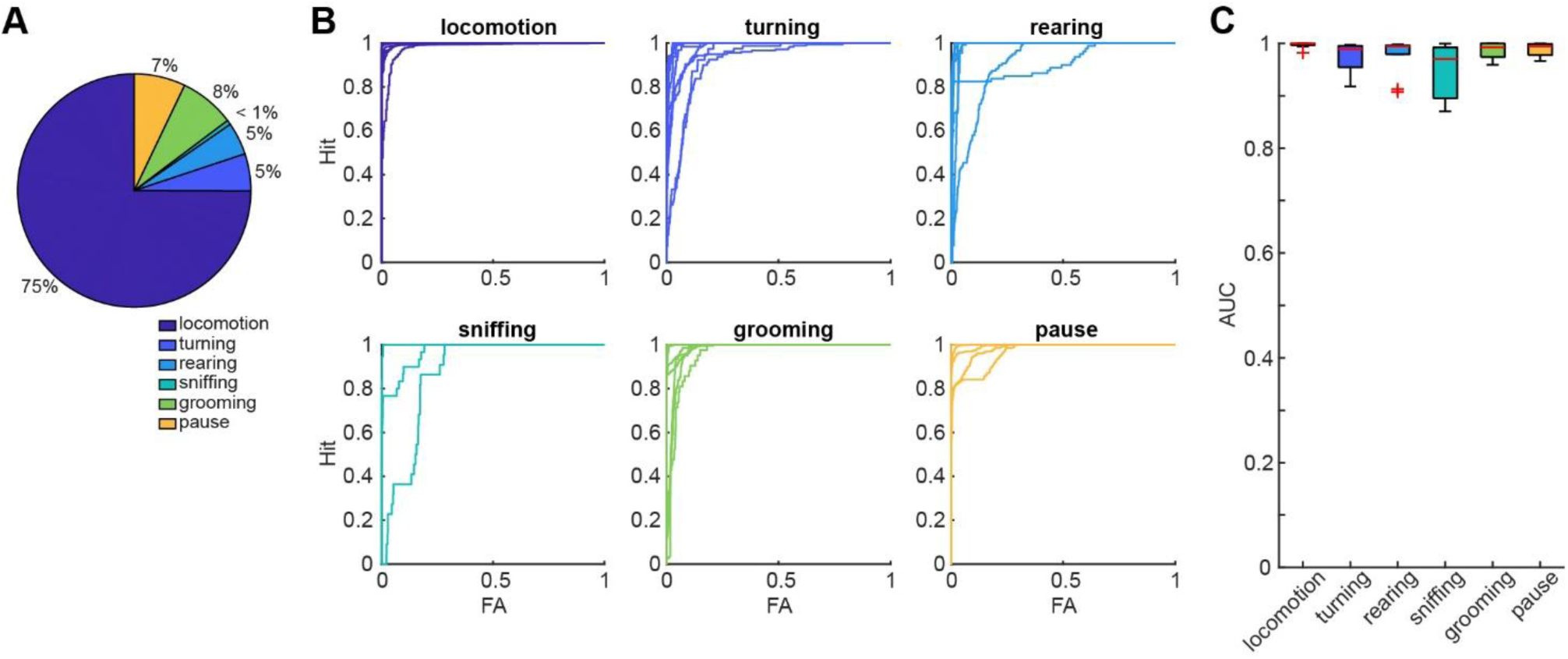
LSTM training data and performance. (**A**) The proportion of each behavioral syllable for LSTM training. (**B**) Receiver operating characteristic curves for each behavioral syllable based on independently labeled 10 videos. FA, false alarm. (**C**) The area under the curve (AUC) values across behavioral syllables. In **Extended Data Figure 3-1**, a range of parameters in the LSTM model were systematically assessed. The evaluation performance of a multiclass support vector machine is shown in **Extended Data Figure 3-2**.

We further evaluated the classifier’s performance by processing additional 10 labeled videos (**Figures 3B and C**). To evaluate evaluation performance, the receiver operating characteristic (ROC) curve (**Figure 3B**) and the area under the curve (AUC) (**Figure 3C**) were calculated for each video and each behavioral syllable. In many cases, the performance was high (>0.95 AUC). As a comparison, a multiclass support vector machine model was also trained. Although the evaluation performance of the trained model was 98.4%, the model failed to generalize the performance to the additional 10 labeled videos (**Extended Data Figure 3-2**). Although the LSTM model’s performance was preserved for those videos, the classification performance for sniffing was not good as other syllables (**Figures 3B and C**). Therefore, we excluded sniffing frames for further analysis (**Figures 5 and 6**). Overall, the trained LSTM classifier provided reliable outcomes across most behavioral syllables.

### Comparisons of SaLSa’s performance with a state-of-the-art model

We compared the performance of SaLSa with that of a state-of-the-art behavioral syllable classification algorithm. Recently, keypoint-MoSeq (kpms) has been introduced and outperforms other models (Weinreb et al., 2023). Therefore, the performance of a kpms model can be a benchmark to evaluate SaLSa. Since we had labelled data from 29 videos, we trained a kpms model with the entire 29 videos. First, we examined how each model classified the labelled data across 6 syllables (**Figures 4A and B**). Consistent with the assessment above, the trained LSTM model provided high classification performance except for sniffing (**Figure 4A**). In the trained kpms model, 15 syllables with one sub-threshold syllable were identified (**Figure 4B**) while all sub-threshold syllables were merged into one syllable. Although some behavioral syllables (locomotion, turning, rearing) were classified into multiple syllables further depending on the direction of the movement, other syllables (sniffing, grooming and pause) tended to be misclassified together. In particular, a significant fraction of grooming behavior was identified as the sub-threshold syllable. We also directly compared the outcomes from two models (**Figure 4C**). The general trend was qualitatively similar to **Figure 4B**. Although locomotion, turning, and rearing were classified into subclasses by the kpms model, other syllables were classified together. To make this trend clearer, syllables from kpms were re-assigned to one of the 6 pre-defined syllables based on the comparison with the labeled data (**Figure 4D**). Then the outcomes of SaLSa and kpms were compared (**Figure 4E**). In this additional analysis, it became clear that kpms misclassified grooming behavior. Thus, although the state-of-the-art model can classify detailed behavioral syllables in a fully automated fashion, SaLSa can reliably classify major behavioral syllables, including grooming.

**Figure 4.**
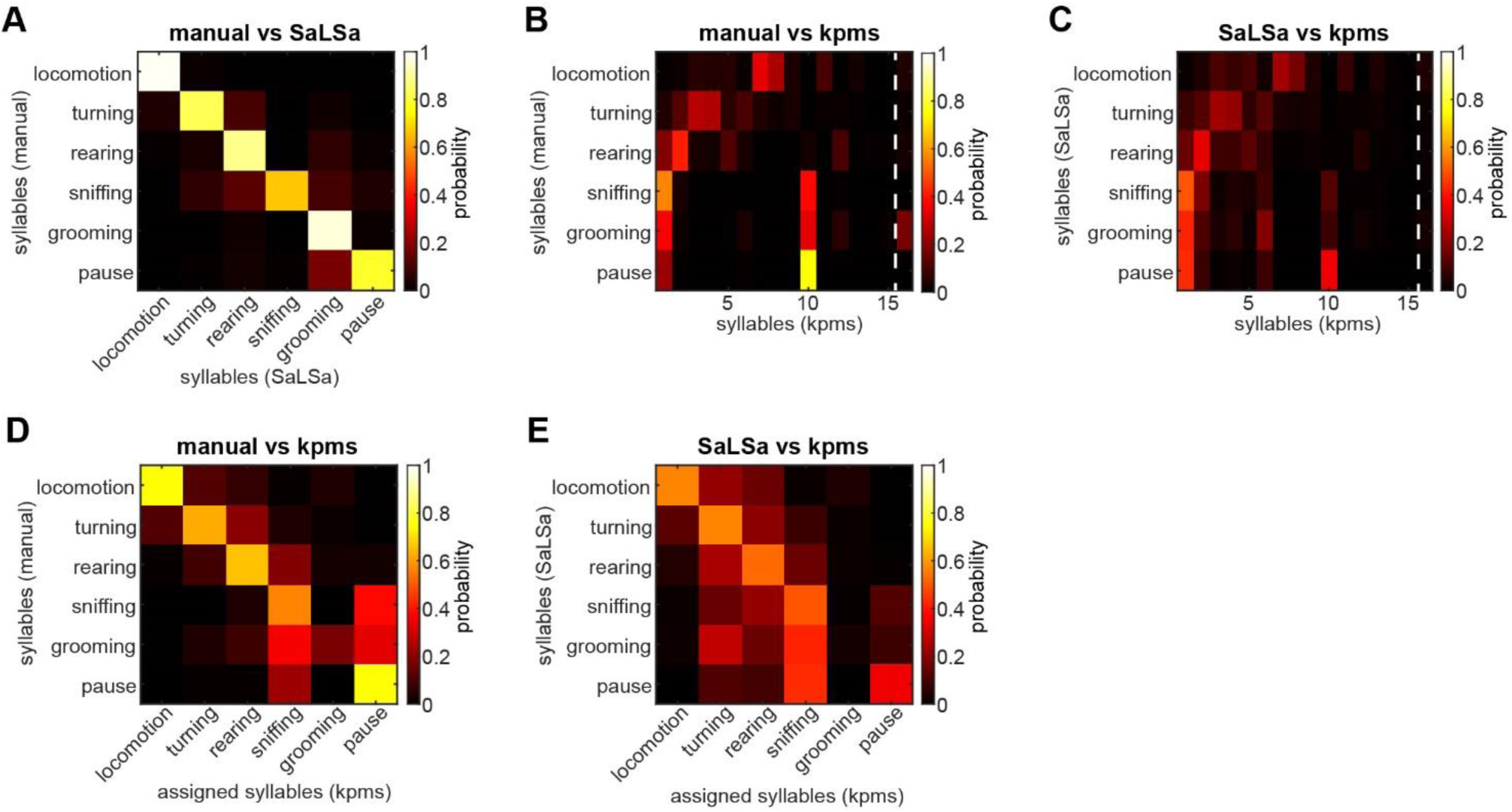
Benchmark testing of SaLSa. (**A**) Confusion matrix between manually labelled data (y-axis) and SaLSa outputs (x-axis). The values indicate what fraction of manually labelled data was classified as each syllable by SaLSa. (**B**) Confusion matrix between manually labelled data and keypoint-MoSeq (kpms) outputs. Syllable 16 is a sub-threshold (0.5% cutoff) class. (**C**) Confusion matrix between outputs from SaLSa and kpms. The values indicate what fraction of frames classified by SaLSa were classified by kpms. (**D**) Confusion matrix between manually labelled data and re- assigned kpms syllables. The original kpms syllables were re-assigned to one of 6 syllables based on comparison to manually labelled data (**B**). (**E**) Confusion matrix between SaLSa output and the re-assigned kpms syllables.

### Quantification of behavioral syllables

Using the trained LSTM classifier, we processed all frames across all 43 videos. To examine if each behavioral syllable has characteristic features, we quantified three simple metrics: speed (**Figure 5A**), turning angle (**Figure 5B**), and compactness (**Figure 5C**). Speed was defined as the nose speed in each behavioral syllable episode. The turning angle was calculated as the cumulative turning angle of the nose-tail base axis in each episode. Compactness was defined as the average pair-wise distance between body parts. We computed the median value of each metric in each video and compared their averages across behavioral syllables (**Figures 5A-C**).

As expected, locomotion was the fastest syllable (F_4, 210_ = 730, *p* < 0.0001, one-way ANOVA with post-hoc Tukey HSD test) whereas grooming and pause were the slowest syllables even compared with rearing (*p* < 0.0005) (**Figure 5A**). The metric of turning angle also provided expected outcomes (**Figure 5B**): turning exhibited the largest turning angle (F_4, 210_ = 456, *p* < 0.01, one-way ANOVA with post-hoc Tukey HSD test) whereas grooming and pause showed a smaller angle than rearing (*p* < 0.0001). Grooming and rearing were the most compact syllables (F_4, 210_ = 75.2, *p* < 0.0001, one-way ANOVA with post-hoc Tukey HSD test) (**Figure 5C**). Because our video monitoring was top-down, rearing typically squeezes their pose in 2-D images. On the other hand, locomotion was the most stretched pose compared to others (*p* < 0.001).

To assess the general structure of animals’ exploratory behaviors in this particular experimental setting, we computed the fraction and average episode duration of each syllable (**Figures 5D and E**). Animals typically spend more time on locomotion and rearing (F_4, 210_ = 129, *p* < 0.0001, one-way ANOVA with post-hoc Tukey HSD test) and less on pause (*p* < 0.0001) (**Figure 5D**). However, while the effect of behavioral syllables on the duration was significant (F_4, 210_ = 66.1, *p* < 0.0001, one-way ANOVA), the duration of grooming was comparable to pause (*p* = 0.18, post-hoc Tukey HSD test) (**Figure 5E**). As expected, turning was the shortest (*p* < 0.0001). Overall, the quantities of each behavioral syllable are consistent with our intuition from behavioral syllables.

### Age-related and sex-specific changes in behavioral syllables of 5xFAD mice

To apply our approach, we examined how abnormalities in 5xFAD mice’s exploratory behavior emerge as they age (**Figure 6**). To this end, we simply compared the fraction of each behavioral syllable as a function of age. The interaction effect between animal groups and age on locomotion was significant (F_3_ = 5.7, *p* < 0.005, ANCOVA): female 5xFAD mice exhibited significantly higher hyper-locomotion compared to female controls (*p* < 0.05, post-hoc Tukey HSD test) (**Figure 6A**). Consistent with this, the interaction effect on pause was also significant (F_3_ = 3.8, *p* < 0.05, ANCOVA) (**Figure 6E**). On the other hand, although the interaction effect on rearing was not significant (F_3_ = 1.85, *p* = 0.155, ANCOVA), the effect of age was significant (F_1_ = 15.4, *p* < 0.0005), meaning that the fraction of rearing decreased with age regardless animal groups (**Figure 6C**). We did not see anysignificant interaction effects on turning and grooming (F_3_ = 2.87, *p* = 0.050 for turning; F_3_ = 1.09, *p* = 0.36 for grooming, ANCOVA) (**Figures 6B and D**). Although hyperactivity of female 5xFAD mice was well described, our approach could dissect detailed behavioral abnormalities.

## Discussion

In the present study, we developed SaLSa, a combination of semi-automatic labeling and LSTM-based classification. The semi-automatic process facilitates the preparation of labeled data for LSTM training whereas LSTM-based classification provides accurate and generalizable behavioral syllable classification. Applying this approach, we found that hyper-locomotion in female 5xFAD mice emerges between 4 and 8 months old whereas other active behaviors, such as rearing, are not affected by genotype or sex. Given its versatility, SaLSa can classify behavioral syllables without an expensive experimental setup.

### Comparisons to other approaches

Over the last decade, a range of approaches has been developed to classify behavioral syllables (Kabra et al., 2013; Pereira et al., 2020; Wiltschko et al., 2020; Dunn et al., 2021; Hsu and Yttri, 2021; Segalin et al., 2021; Jia et al., 2022; Luxem et al., 2022; Harris et al., 2023; Luxem et al., 2023; Weinreb et al., 2023). Since these approaches including SaLSa are applied after body parts detection, they can be applied to videos taken in relatively dark environments as long as body parts are detected reliably. SaLSa is the first approach to utilize LSTM for this purpose, to the best of our knowledge. Deep learning-based approaches have increasingly been adopted for this purpose (Marks et al., 2022; Harris et al., 2023). Although convolutional deep learning models are powerful to classify and segment images, they lack an intrinsic mechanism to hold contextual information. Recurrent neural networks are suitable to process time series data, including behavioral tracking data (Luxem et al., 2022). One advantage of LSTM over conventional recurrent networks is that it can learn long-term dependencies on the data (Hochreiter and Schmidhuber, 1997). Because the brain can deal with several orders of magnitudes of time depending on its computational goals (Issa et al., 2020), adopting LSTMs is an extension of ongoing efforts to characterize natural behaviors comprehensively. Another advantage of LSTM is its generalizability like many other deep learning approaches. Once it has been trained, newly acquired videos can be processed by simply extracting features as we demonstrated.

On the other hand, LSTM requires a large amount of labeled data for training. To mitigate this issue, we adopted semi-automatic labeling. By extracting features unbiasedly, behavioral syllable candidates were automatically identified. Compared to an approach that creates snippets randomly (such as MuViLab), our approach reduces manual curation time. In the present study, a 20-minute video took several minutes. This allows labeling a number of videos easily to prepare training data. Thus, our approach is unique in the sense that it combines both unsupervised methods and a supervised LSTM model to classify behavioral syllables.

In a broader context, SaLSa takes a top-down approach where pre-defined behavioral syllables are identified semi-automatically and classified by a deep learning model. In the future, it may be interesting to integrate a fully automated bottom-up model with an LSTM classifier.

### Behavioral abnormalities in 5xFAD

As an application of SaLSa, we investigated age- and sex-specific changes in the exploratory behaviors of 5xFAD mice. It has been well documented that female 5xFAD mice exhibit hyperactivity (Oblak et al., 2021). Our approach demonstrates that hyperactivity consists of hyper-locomotion whereas other active behaviors, such as turning and rearing, are similar across animal groups. In particular, rearing decreases with age regardless of genotype and sex, which has not been documented in this mouse model before.

The underlying mechanisms of sex-specific hyper-locomotion in 5xFAD mice are unknown. In this mouse model, amyloid plaques can be seen in the hippocampus (primarily the subiculum) as early as 2 months old and the pathology appears across brain regions as they age (Oakley et al., 2006; Oblak et al., 2021). Sex differences in amyloid pathology can also be apparent in the cortical subplate even at 3 months old and seen across multiple brain regions at 4 months old (Oblak et al., 2021). Consistent with this sex-specific pathological progression, a transcriptomic analysis also revealed sex differences in a wide range of molecular pathways (Oblak et al., 2021). In the future, it would be crucial to determine how these sex-specific pathological features link to dysfunctions in neural circuit activity, which lead to age-related, sex-specific hyper-locomotion. Despite this challenge, given the simplicity of our experimental setup, similar approaches can be applied to other animal models.

### Limitations of the study

Our study has at least four limitations: first, our pre-set behavioral syllables were limited. We are also aware that some animals jumped or exhibited complex behaviors, such as mixed turning and rearing behaviors. This will require more labeled data or additional post-hoc analysis. For example, based on the predicted score from a classifier, such complex behavioral syllables may be defined. Additionally, sniffing behaviors could not be identified and classified accurately. This could be partly because of the limited resolution of our camera and the accuracy of body-part tracking. Increasing the resolution and adding extra body parts to track may improve this aspect.

Second, the model must be re-trained when videos are taken at different frame rates or from a different experimental setup. Because our temporal features assume a certain frame rate (25 fps), a change in frame rate leads to a change in the number of features. To deal with this issue, re-sampling can be considered.

Third, while the present study replicated the results (i.e., sex-specific hyperactivity) of the recent comprehensive analysis in 5xFAD mice with the detailed classification of behavioral syllables, the estrous cycle might be a potential confounding factor even though a recent study could not find this to be the case (Levy et al., 2023). Increasing the sampling size by monitoring the estrous cycle will address this issue in the future.

Finally, as widely appreciated, deep learning models are challenging to interpret. In this study, it is not straightforward to determine what spatiotemporal features contribute to classification. Several approaches may be considered, such as Local Interpretable Model-Agnostic Explanations (LIME) and visualization.

### Conclusions

Behavioral syllable classification is important. In the present study, we developed a combinatory approach to semi-automatic labeling and LSTM-based classification, called SaLSa. Our approach can assist manual curation to prepare labeled data semi-automatically. The LSTM classifier reliably classified behavioral syllables in new datasets, which were not used for training. It also provides comparable performance with the state-of-the-art model. Thus, our approach adds a versatile tool for behavioral syllable classification. Combining other advanced technologies, SaLSa facilitates the effort to better understand the neural basis of complex behavior.

## Acknowledgements

We thank Abigail Hatcher Davies for her technical assistance. This work was supported by MRC (MR/V033964/1) and Horizon2020-ICT (DEEPER, 101016787) to SS.

## Extended Data Figures

**Extended Data Figure 3-1.**
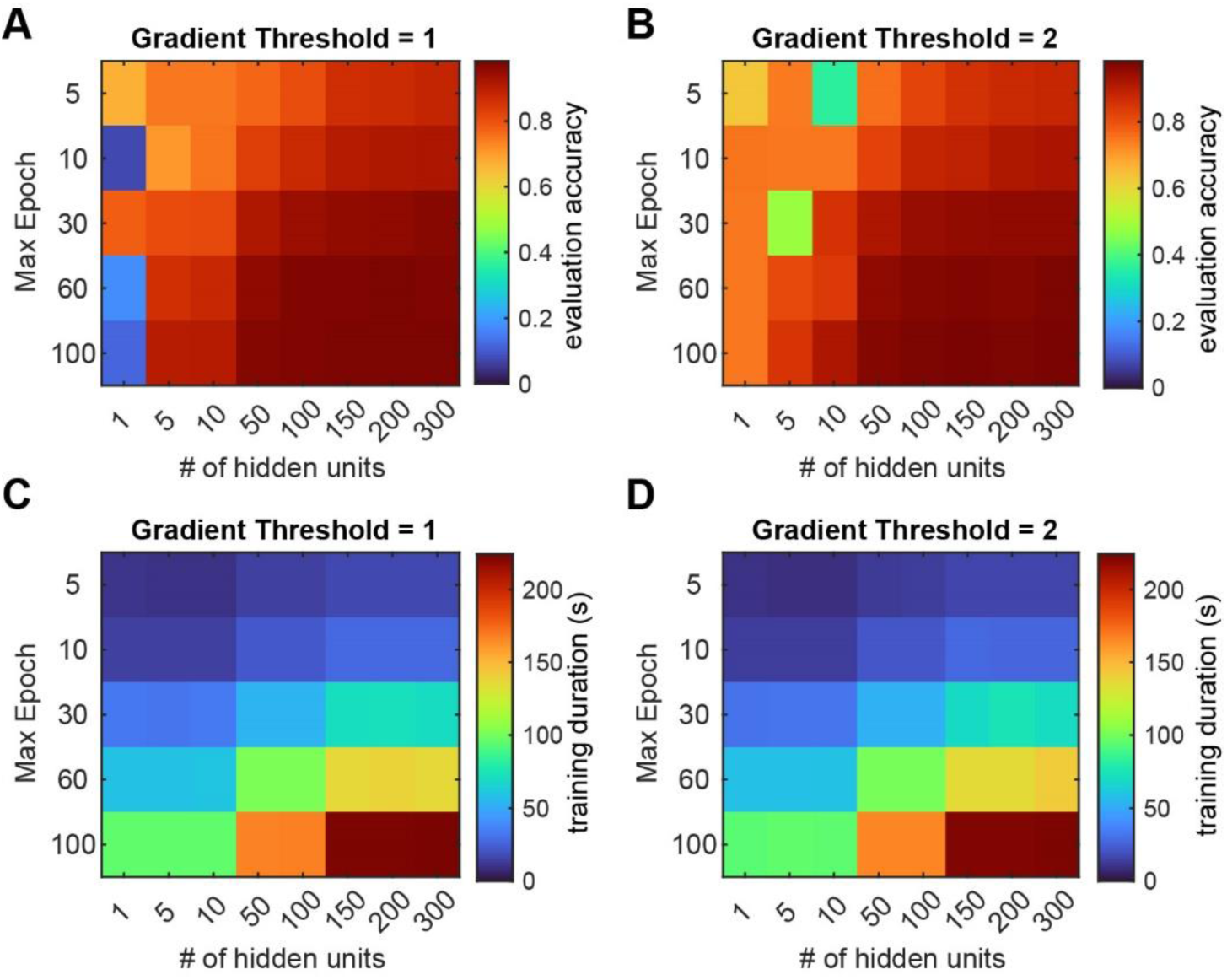
Systematic comparison of three parameters for evaluation accuracy and training duration. (**A and B**) The evaluation accuracy of LSTM models with variable maximum numbers of epochs and hidden units. The gradient threshold was set at 1 in (**A**) and 2 in (**B**). (**C and D**) Training duration across conditions.

**Extended Data Figure 3-2.**
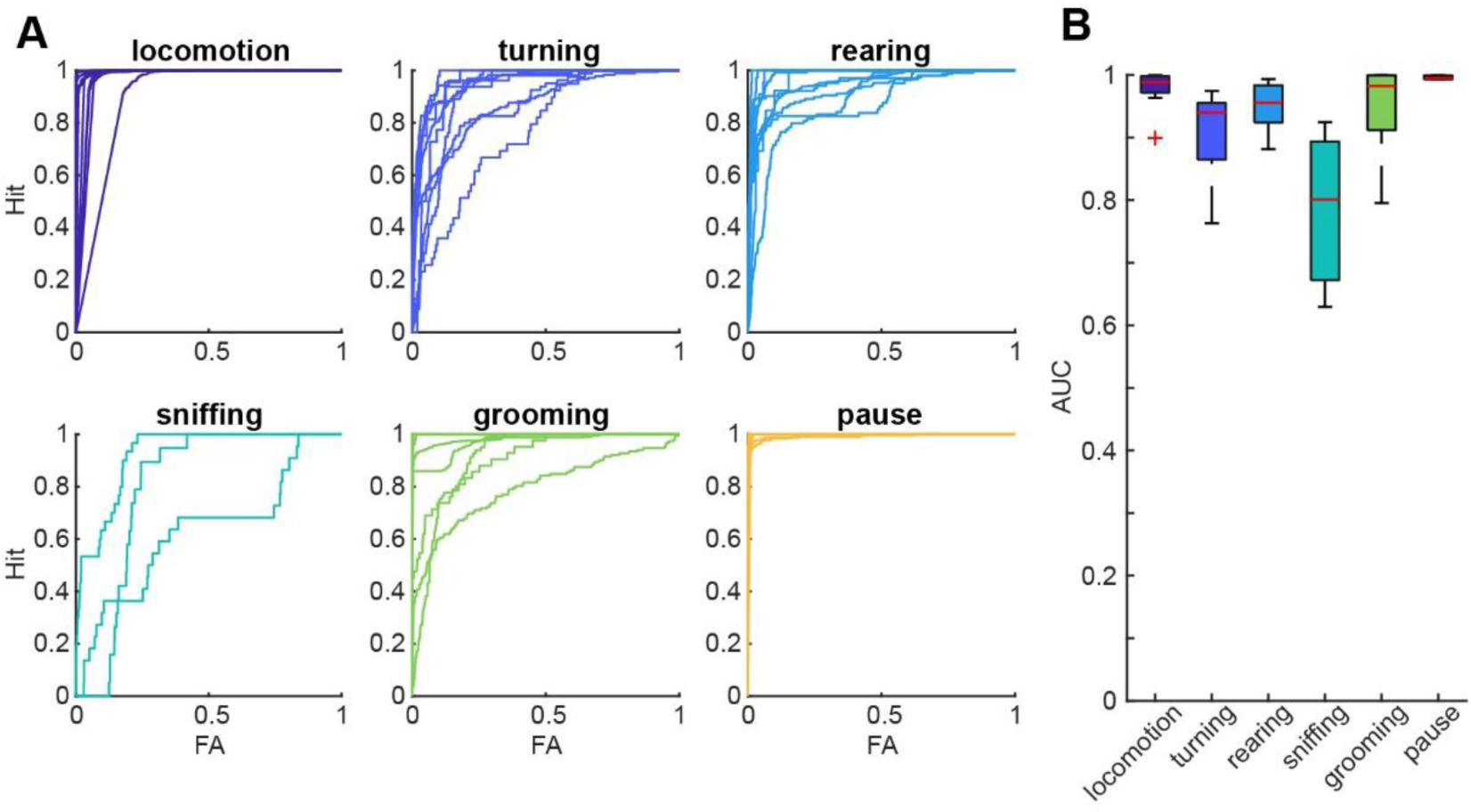
Performance of a multiclass support vector machine. (**A**) Receiver operating characteristic curves for each behavioral syllable based on independently labeled 10 videos. FA, false alarm. (**B**) The area under the curve (AUC) values across behavioral syllables.

